# Principles of tactile search over the body

**DOI:** 10.1101/839084

**Authors:** Elizabeth J. Halfen, John F. Magnotti, Md. Shoaibur Rahman, Jeffrey M. Yau

**Author notes:** Corresponding author: Jeffrey M. Yau One Baylor Plaza, T111 Houston, TX 77030.

## Abstract

Although we experience complex patterns over our entire body, how we selectively perceive multi-site touch over our bodies remains poorly understood. Here, we characterized tactile search behavior over the body using a tactile analog of the classic visual search task. Participants judged whether a target stimulus (e.g., 10-Hz vibration) was present or absent on the upper or lower limbs. When present, the target stimulus could occur alone or with distractor stimuli (e.g., 30-Hz vibrations) on other body locations. We varied the number and spatial configurations of the distractors as well as the target and distractor frequencies and measured the impact of these factors on search response times. First, we found that response times were faster on target-present trials compared to target-absent trials. Second, response times increased with the number of stimulated sites, suggesting a serial search process. Third, search performance differed depending on stimulus frequencies. This frequency-dependent behavior may be related to perceptual grouping effects based on timing cues. We constructed models to explore how the locations of the tactile cues influenced search behavior. Our modeling results reveal that, in isolation, cues on the index fingers make relatively greater contributions to search performance compared to stimulation experienced on other body sites. Additionally, co-stimulation of sites within the same limb or simply on the same body side preferentially influence search behavior. Our collective findings identify some principles of attentional search that are common to vision and touch, but others that highlight key differences that may be unique to body-based spatial perception.

**New & Noteworthy:** Little is known about how we selectively experience multi-site touch over the body. Using a tactile analog of the classic visual search paradigm, we show that tactile search behavior for flutter cues is generally consistent with a serial search process. Modeling results reveal the preferential contributions of index finger stimulation and two-site interactions involving ipsilateral and within-limb patterns. Our results offer initial evidence for spatial and temporal principles underlying tactile search behavior over the body.

## Introduction

Our sensory environments are often complex and cluttered. Because our capacity to process simultaneously experienced sensory inputs is limited, efficient processing of complex environments requires mechanisms for identifying salient and meaningful signals while filtering out noise and distractors. Spatial attention enables us to allocate limited neural resources to behaviorally relevant signals in our surroundings. While much is known regarding the selection and filtering of signals in cluttered visual and acoustic environments, the selective processing of simultaneously experienced tactile cues remains poorly understood, despite the fact that mechanoreceptors tile our entire body surface and we are constantly experiencing dynamic tactile inputs at multiple locations over our body.

Within the field of visual spatial attention, the visual search procedure has been widely used to investigate how we allocate attention (Eckstein 2011; Wolfe and Horowitz 2017). In visual search, subjects are tasked with reporting the presence or absence of a target stimulus amongst a varying number of distractor stimuli. Response times on visual search tasks can depend strongly on the target and distractor features as well as the number of stimuli presented in the visual field (i.e., set size). The relationship between response times and set size has long been used as an index of search efficiency (Duncan and Humphreys 1989; Treisman and Gelade 1980). Search behavior is considered efficient when response times remain unchanged as set size increases; efficient search is interpreted as evidence for parallel processing of the sensory cues. Conversely, search behavior is considered inefficient when response times increase with set size increases; inefficient search is interpreted as evidence for a serial search process where each stimulus in the display is sequentially evaluated. By identifying features and contexts associated with serial and parallel search, “guiding principles” have been established that provide a framework for understanding visual search behavior (Wolfe and Horowitz 2017) and its potential neurophysiological basis (Bichot et al. 2005; Cohen et al. 2009).

Touch and vision are similar in a number of ways, including the type of spatial information represented by these senses and the manner in which attention modulates this information (Hsiao et al. 1993). Indeed, we perceive similar features with touch and vision (Pei and Bensmaia 2014; Phillips et al. 1983; Yau et al. 2016), and the somatosensory and visual cortical systems employ analogous coding principles to represent these features (Bensmaia et al. 2008; Pei et al. 2011; Yau et al. 2009). Tactile spatial attention has also been shown to be similar to visuospatial attention (Gallace and Spence 2011; Johansen-berg and Lloyd 2000). For example, spatial cueing in a Posner-like paradigm reduces response times to tactile stimuli as it does with vision, and cueing facilitates processing, even in crossmodal contexts (Butter et al. 1989; Whang et al. 1991). Additionally, touch and vision both exhibit inhibition of return: processing of a previously attended location in space is inhibited for a short period of time after attention is allocated there (Cohen et al. 2005; Poliakoff et al. 2002). Given the many similarities between visual and tactile spatial processing, we reasoned, like others (Lederman and Klatzky 1997; Overvliet et al. 2007; Toet et al. 2008), that the principles underlying tactile attentional search should be relatable to the guiding principles of visual search.

Here, we characterized tactile search behavior using a full-body search paradigm analogous to the classic visual search paradigm. Our design differed from previous efforts to investigate tactile search, which primarily focused on attentional selection over the hands (Lederman et al. 1988; Lederman and Klatzky 1997; Overvliet et al. 2007) or over a single expansive body part like the torso (Toet et al. 2008). In contrast, we delivered target and distractors cues (10-Hz and 30-Hz “flutter” trains) to 8 sites covering distal and proximal locations on the arms and legs on both sides of the body. By distributing tactile cues over multiple body parts and limbs, we hoped to minimize the potential confounding effects associated with physical interactions between tactile cues experienced at nearby locations on the body or interaction effects like masking or crowding (Gilson 1969; Levin and Benton 1973; Mahar and Mackenzie 1993; Sherrick 1964), which may be more pronounced with tactile stimulation on the hands (Craig 1985; D’Amour and Harris 2014a; Sherrick 1964) or on a single body part (Levin and Benton 1973; Mahar and Mackenzie 1993). We characterized the effect of target presence, set size, and target frequency on search behavior. We also manipulated the phase of the distractor cues to explore the role of timing cues in stimulus grouping effects. Lastly, we constructed linear models to infer the influence of individual body sites and pairs of co-stimulated body-sites on response times. Taken together, our results establish a preliminary framework for understanding spatial search behavior over the body.

## Materials and Methods

### Participants

We recruited a total of 16 subjects (9f; mean age ± standard deviation: 27.9 ± 5.3 years). We were unable to establish reliable iso-intensity values for the 10- and 30-Hz test stimuli across sites (see below) in 4 subjects, so these individuals were not tested in the main experiments. Of the remaining 12 subjects, 10 were right-handed according to the Edinburgh Handedness Inventory (Oldfield 1971) and all subjects reported normal somatosensory functions. Testing procedures were performed in compliance with the policies and procedures of the Baylor College of Medicine Institutional Review Board. All participants provided informed written consent and were paid for their participation or declined compensation.

### Vibrotactile stimulation

Tactor stimulation was delivered to 8 body locations using electromechanical tactors (Engineering Acoustics, Inc., EAI) (Convento et al. 2018; Rahman and Yau 2019) (Fig. 1A). Tactile stimuli were generated and controlled using the EAI TDK for MATLAB (2018a, MathWorks) on a PC laptop (model GL503VS; Windows 10 Home, 2.8GHz Core i7, 16GB RAM). Using the EAI software and hardware, 10- and 30-Hz stimulus trains were generated by pulse width modulation (pulse width: 10ms; carrier frequency: 200Hz). Experiment timing and visual cues were controlled using Psychtoolbox-3 (Kleiner et al. 2007). Subjects listened to white noise (Audio-Technica ATH-ANC23 QuietPoint Active Noise-Cancelling In-Ear Headphones) to mask any sounds produced by the tactors.

**Figure 1.**
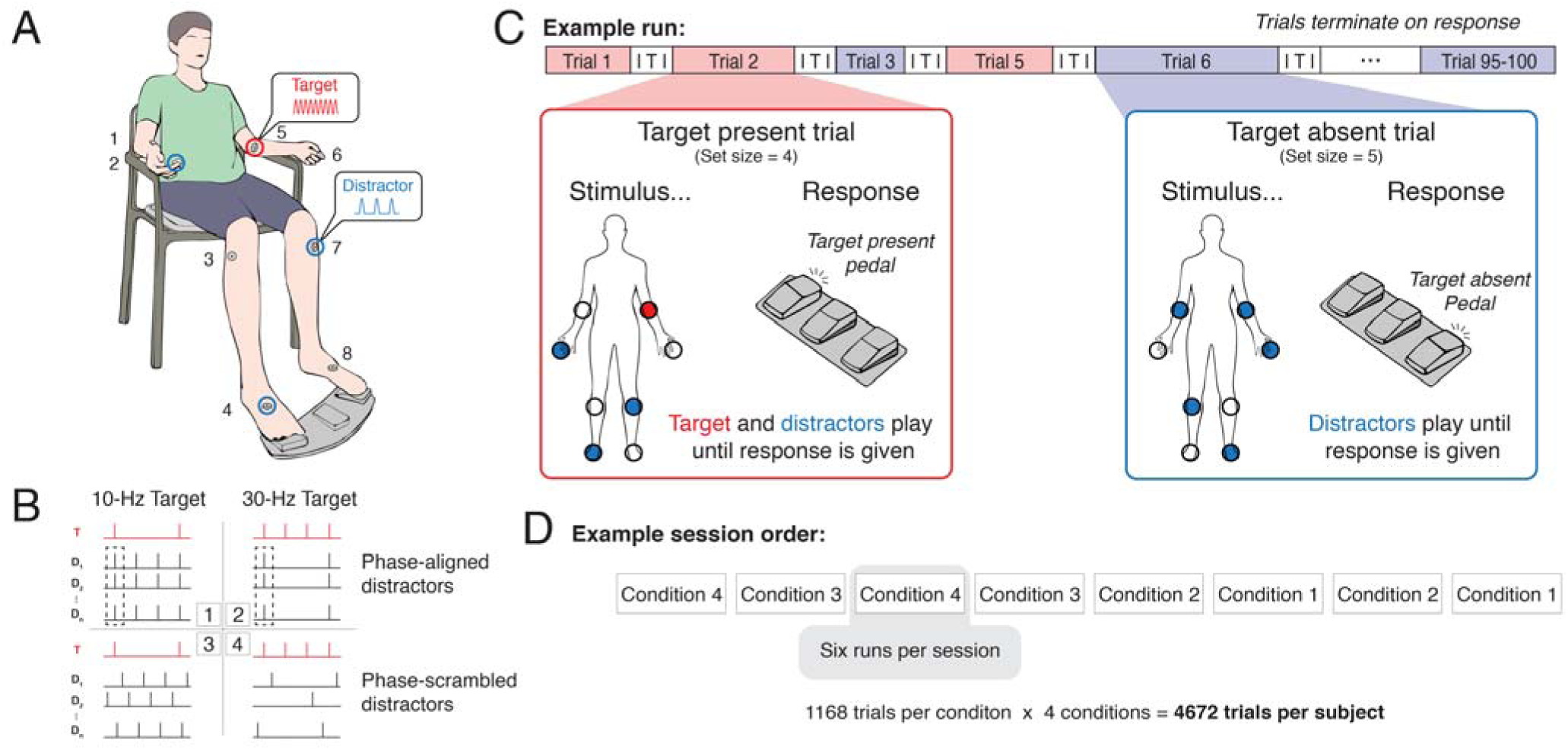
Experimental design. (***A***) Subjects were seated in a chair with their arms resting at their sides. The 8 numbered sites indicate where the tactors were affixed to the subject’s body. Subjects rested their feet on the left and right pedals of a three-pedal response device. On each trial subjects reported the presence or absence of a target stimulus (red) in the context of concurrent distractor stimulation (blue). (***B***) Schematics of tactile stimulus trains showing 4 conditions tested (inset boxes). Red trains depict target stimulus and black trains depict distractor trains under 2 target frequency conditions (columns) and 2 distractor phase conditions (rows). Black dashed boxes highlight synchronized distractors in the phase-aligned condition. (***C***) An example run comprising target-present (red) and target-absent (blue) trials. Each trial terminates on subject response. Inter-trial interval (ITI) varied between 2-2.5s. Each run consisted of 95-100 trials. (***D***) Subjects completed 8 sessions of tactile search. Phase-aligned distractors and phase-scrambled distractors were tested in 4 sessions each and stimulus frequency condition was alternated over sessions (Conditions from *B*). The order of both were counterbalanced across subjects. Each session consisted of 6 runs. Each subject was tested with 4,672 total trials.

Different EAI tactors were used depending on the stimulation site. On the left and right sides, C-F tactors (0.5 cm diameter contact) were used on the index finger, C-3 tactors (0.8 cm diameter) on the forearm, C-2 tactors (0.8 cm diameter) on the lower leg, and C-2 tactors on the foot. Index finger tactors were centered on the distal fingerpad. Forearm tactors were positioned medially near the elbow, 20% down the length of the ulna. To position the lower leg tactor, we first measured the distance from the middle of the patella to the middle of the talus (i.e., the bend of the ankle) and placed the tactor 20% along this distance from the knee and 2 cm laterally. The foot tactor was positioned in the middle of the dorsal surface of the foot. When fastened to the feet, knees, or forearms, tactors were taped to the body (Transpore tape, 3M). When fastened to the finger, the tactors were fixed using an elastic fabric band with Velcro (Convento et al. 2018). Tactor positions were established during each participant’s first test session and maintained over all sessions.

### Experimental Design

#### General overview

Each participant first completed 2-3 sessions of intensity matching experiments prior to beginning the tactile search experiments. They then completed 2 tactile search experiments (phase-aligned experiment and phase-scrambled experiment), each of which consisted of 4 sessions. Participants completed a full experiment before beginning the other: half of the participants began with the phase-aligned experiment while the other half began with the phase-scrambled experiment. Thus, each participant completed a total of 10 or 11 sessions.

#### Intensity matching

To account for sensitivity differences across different body locations for each participant, we established the amplitudes of the 10- and 30-Hz stimuli at each stimulation site that were perceived to be equally intense as a 10-Hz reference stimulus delivered to the right knee at a fixed amplitude (70 arbitrary units (a.u.) on the EAI Universal Controller). This reference stimulus level was chosen as it was clearly suprathreshold, but also well below amplitudes that would be perceived as uncomfortable. Subjects first matched the perceived intensity of 10-Hz stimuli across body sites, by way of seven 2-site comparisons. The order of these across-site comparisons was randomized; however, matching between homologous body parts (i.e., left to right hand) was prioritized. After completing across-site matching, subjects then matched the perceived intensity of 10- and 30-Hz stimuli at each body site.

In this experiment and all others, subjects sat comfortably in a chair with their hands supinated (Fig. 1A), centered in front of the monitor on which instructions were delivered. Iso-intensity values were determined using a two-interval, two-alternative forced choice paradigm and an adaptive tracking procedure (QUEST, Watson and Pelli 1983). On a given run, subjects performed an amplitude discrimination task involving same-frequency stimuli experienced at 2 different locations or different-frequency stimuli experienced at the same location. A text prompt on the screen indicated which body site(s) were to be compared. On each trial, participants experienced two stimuli sequentially (stimulus duration: 1.5s; inter-stimulus interval: 0.5s) before verbally reporting which stimulus was perceived to be higher in intensity. The experimenter recorded each response with a key press. On each trial, one stimulus was always the reference cue (i.e., the 10-Hz right knee stimulus or another stimulus previously equated to the 10-Hz right knee stimulus) and the amplitude of the other (comparison) stimulus was varied according to the adaptive algorithm. Each run comprised 2 interleaved adaptive sequences, each consisting of 30 trials, which differed according to the initial amplitudes of the comparison stimuli. Each run lasted 6-8 minutes. The final estimates from the 2 adaptive sequences were averaged for the iso-intensity value. If the 2 adaptive sequences did not yield consistent results, defined as sequence convergence to values within 40% of the mean result, the run was repeated. Subjects who were unable to achieve consistent iso-intensity values were excluded from further testing. Four subjects were excluded, yielding a total of 12 subjects with iso-intensity values for 10- and 30-Hz stimuli over 8 body sites.

#### Tactile search with phase-aligned distractors

The tactile search experiment is analogous to the classic visual search task (Treisman and Gelade 1980). On each trial, subjects reported whether a tactile stimulus of a specified (target) frequency was detected at any of the 8 potential stimulation sites on the body (Fig. 1A). Subjects made “Target Present” and “Target Absent” responses using different foot pedals (Lemo 1640835 PC USB Foot Control Keyboard Action Switch Pedal HID). Subjects were instructed to respond as quickly as possible without needing to localize the target before responding. During each trial, the stimuli stopped once a response was given or 10s had elapsed without a response (0.25% of trials). No response feedback was provided. The assignment of “Target Present” and “Target Absent” responses to the left and right foot pedals was counterbalanced across subjects.

In separate sessions, the target frequency was either 10 Hz (Condition 1; with 30-Hz distractors) or 30 Hz (Condition 2; with 10-Hz distractors). Each session comprised 6 runs of trials. Subjects were instructed to maintain visual fixation on a central cue throughout a run. Three participants performed the experiment without the constraint to maintain visual fixation, but their data were similar to that of the other participants. The target stimulus was present on 70% of the trials (Fig. 1C). The remaining 30% of trials only included distractor stimulation (Fig. 1C). When multiple distractors were delivered in parallel during a trial, the pulses were delivered synchronously (Fig. 1B). Trials were additionally classified according to set size – the number of stimulated sites – which ranged from 1 to 8. Set sizes were sampled nonuniformly (Table 1), but all set sizes were tested at least 24 times within each subject. For target-present trials, we tested all stimulus patterns of set sizes 1-3 and 7-8 (ranging from 1-5 repetitions of a given pattern); however, we tested only a random subset of possible stimulus patterns classified by set sizes 4-6 (each with a single repetition). For target-absent trials, stimulus patterns were sampled randomly with repeat from all possible target-absent combinations for each set size.

**Table 1.**
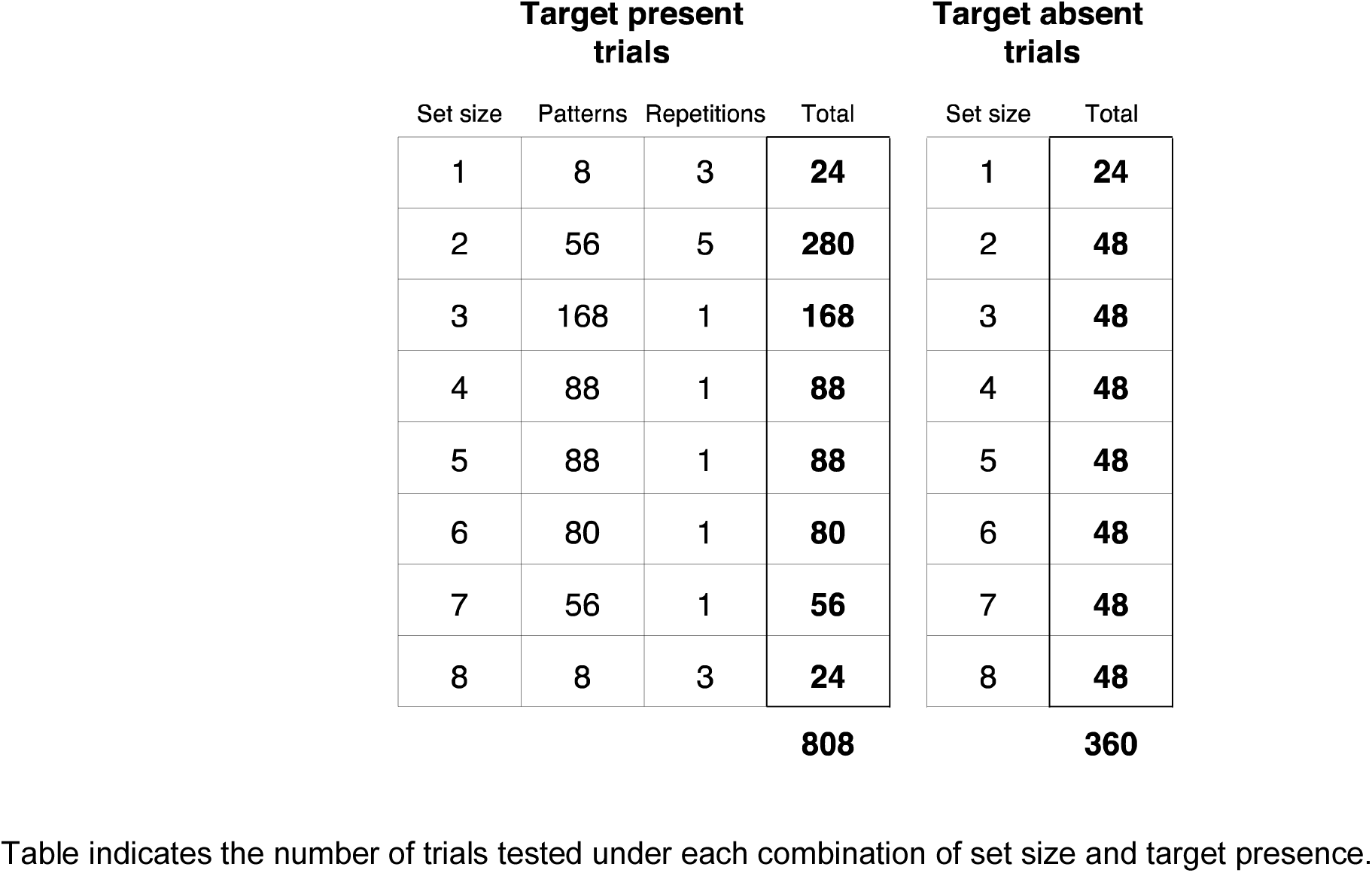
Trial distribution over all combinations of set size and target presence.

Stimulus patterns and order were separately determined for each subject. Individual runs were balanced with approximately the same number of trials for each set size and target presence combination. At the beginning of each session, subjects participated in a short practice run to familiarize them with the target frequency. Each session consisted of 6 runs (95-100 trials per run). Subject were allowed to rest for 1-2 minutes between each run. Conditions 1 and 2 were each tested twice over the 4 sessions (Fig. 1D), the order of which was counterbalanced across subjects (within-Condition inter-session interval: 3 ± 2 days). Across the experiment, we acquired behavioral responses from 1168 trials for each condition from each participant.

#### Tactile search with phase-scrambled distractors

This experiment employed the same design as described above except that co-occurring distractors were delivered out of phase (Fig. 1B; Conditions 3 and 4 for the 10- and 30-Hz targets, respectively). Phase delays were randomly selected without replacement from binned intervals of the period of the distractor frequency. This design ensured that pairs of distractors were never synchronized, allowing us to test the hypothesis that timing cues (i.e., phase-alignment) contribute to the perceptual grouping of stimulus trains experienced across the body. Six subjects completed the tactile search experiment using phase-scrambled distractors before participating in the tactile search experiment using phase-aligned distractors.

### Statistical analysis

The primary analysis goal was to determine how response time (RT) depended on set size, target presence, and stimulus frequency. We report response times rounded to the nearest millisecond. Analyses were performed using MATLAB (R2019a) and RStudio (R version 3.5.3). We modeled how RT changed across conditions using linear mixed effects modeling as implemented in the lme4 package (Bates et al. 2015). We fit separate models to the data from Experiment 1 (distractors phase-aligned with target) and Experiment 2 (phase-scrambled distractors). Each model included fixed effects of Set Size (1 to 8), Target Status (Present vs. Absent), Target Frequency (10Hz vs. 30Hz), and their 2- and 3-way interactions. Subject identification was included as a random effect in each model; no other random effects were included because the model optimization did not converge. Models were optimized using log-likelihood (option REML=FALSE). We used the lmerTest package (Kuznetsova et al. 2017) to provide tests of significance for all fixed effects using the Satterthwaite method for degrees of freedom calculation. The reported degrees of freedom are rounded to the nearest integer.

#### Modeling the response time contributions of individual body sites

To evaluate how tactile stimulation at each stimulated site independently affected search performance over the body, we constructed a linear model to predict response times for each stimulus pattern (including target present and target absent trials across all set sizes), using correct trials only. The model assumed that response time is described as the weighted sum of the contributions of the stimulated body sites:

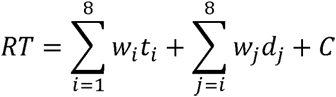

where *w_i_* is the response time effect of a target stimulus *t* at the *i*^th^ site, *w_j_* is the contribution of a distractor stimulus *d* at the *j*^th^ site (note *i*≠*j*), and *C* is a constant that accounts for the overall magnitude of the response times. For each subject, we used least squares regression to obtain the best fitting values for all *w_i_* and *w_j_*. Different parameters were fitted for the two target frequencies under the phase-aligned and phase-scrambled conditions, yielding 4 distinct sets of coefficient estimates.

#### Modeling 2-site interaction effects on response time

To test if interactions between pairs of body sites accounted for response time variance unexplained by the linear model, we constructed a model that comprised not only linear target and distractor terms (as in the previous section), but also interaction terms that modeled the response time contributions of pairs of simultaneously-stimulated body sites:

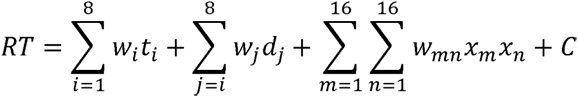

where *w_mn_* is the weight for the co-occurrence of the *m* and *n* conditions, each describing a unique stimulus x site condition (target or distractor at one of 8 sites). Because a target and distractor cannot occur at the same site, and because two targets cannot occur in the same trial, only 84 two-site terms were possible. With one-site terms, valid two-site terms, and the constant included, the most comprehensive model comprised 101 terms.

Rather than assuming this maximally complex model and that all two-site terms were statistically meaningful, we quantified the relative contributions of each two-site term and objectively determined whether their inclusion in the model was justified using a modified orthogonal matching pursuit algorithm (Mallat and Zhang 1993) and a rigorous model selection process. We again fitted separate models to each subject’s data for each target frequency condition and distractor-phase condition. Matching pursuit is an iterative sparse approximation algorithm: On each iteration, the algorithm identifies the term from a dictionary (here, all of the valid two-site terms) that best accounts for the explainable variance remaining from the previous iteration. This term’s contribution is then removed from the data and the process is repeated until all terms have been selected from the dictionary or none of the remaining terms describe any of the remaining data variance. This procedure thus yields a sequence of two-site terms ordered by their relative contributions to response time.

Because this procedure is deterministic and sensitive to noise, we adopted a randomization approach to establish term sequence (Thakur et al. 2012). Rather than selecting the term with the largest projection onto the signal on each iteration, we randomly selected a term from those that had projections >75% of the top projection in fitting the full model. We then repeated this entire process 10,000 times, which resulted in 10,000 ordered term sequences. To determine the ultimate term sequence for a given model – the model most consistent and robust to noise during fitting – we set the two-site term for a given sequence position (e.g., the 4^th^ two-site term) as the term that occurred most frequently at that sequence position over the 10,000 sequences, without replacement.

To determine how many two-site terms should be included in the final model, we performed model selection using Bayesian Information Criterion (BIC) (Schwarz 1978). The full set of alternative models consisted of 85 models that varied only according to the number of two-site terms with the term sequences determined by the randomized matching pursuit procedure. All potential models included the 16 one-site terms as well as the constant. We computed BIC for each model:

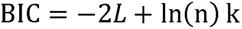

where *L* is the log likelihood of our observed response time data given the predictions of a model with *k* parameters and *n* is the number of unique trial types. To calculate the likelihood of the data given a fitted model, we used separate Gaussian distributions for the response times associated with target-absent and target-present trial types. The preferred model was the model associated with the lowest BIC value.

In estimating the single-site and 2-site models we chose to fit per subject rather than constructing a single linear mixed-effects (LME) model with stimulation-site and site-interaction terms. The hallmark of the LME approach is the partial pooling of data across subjects (the random effect term); however, this partial pooling is only optimal if we assume that all models are (Gaussian) samples from a single “true” model. Instead, fitting each individual separately and post-hoc comparing the results allows us to directly assess individual variation. Looking at single-subject data confirms the validity of our approach in this instance, as the number of terms selected for different subjects varied greatly (see below).

#### Model fitting and performance

To quantify model similarity between conditions and subjects, we calculated Hamming distance (*D_H_*) (Hamming 1950):

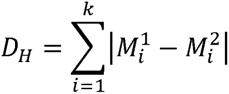

where *M_i_*^1^ and *M_i_*^2^ indicate the presence or absence of the *i*^th^ term of the models *M^1^* and *M^2^*, each of which is a binarized vector of length k (the total possible two-site terms). Lower *D_H_* values indicate greater similarity between the models. We determined the similarity of 10-Hz and 30-Hz models for each subject by concatenating the models over distractor phase. We evaluated whether the mean within-subject *D_H_* was significantly lower than expected by chance (p < 0.05) by generating null distributions of 1000 distance values computed from randomly shuffled *M^1^* and *M^2^* vectors for each subject and drawing from these distributions to generate a (group-level) null distribution of 1000 mean distance values against which we compared the observed mean *D_H_*. We also evaluated model similarity across subjects by calculating *D_H_* between the models of pairs of subjects, concatenated over target frequency and distractor phase. The observed mean between-subjects *D_H_* was compared to a null distribution of mean distance values computed from permuted data.

### Data and Code Availability

The behavioral data and analysis code are available at https://github.com/YauLab/TactileSearch.

## Results

### Performance accuracy on target-present and target-absent trials

Although we focused our analyses on response times, we first established how well participants performed the tactile search task. Participants achieved high performance levels with both phase-aligned and phase-scrambled distractors, with subtle differences that depended on the target frequency or whether the target was present or absent. With phase-aligned distractors (Fig. 2A), performance tended to be higher on target-absent trials (percent correct; 10-Hz target: 98.03 ± 0.44%; 30-Hz target: 96.74 ± 0.90%) compared to target-present trials (10-Hz target: 73.37 ± 4.15%; 30-Hz target: 89.63 ± 2.31%). A similar pattern described performance with phase-scrambled distractors (Fig. 2B) as we observed better performance on target-absent trials (10-Hz target: 97.15 ±1.33%; 30-Hz target: 94.19 ± 2.73%) compared to target-present trials (10-Hz target: 73.60 ± 4.88%; 30-Hz target: 91.01 ± 2.02%. We restricted all subsequent analyses of response times to those measured in correct trials.

**Figure 2.**
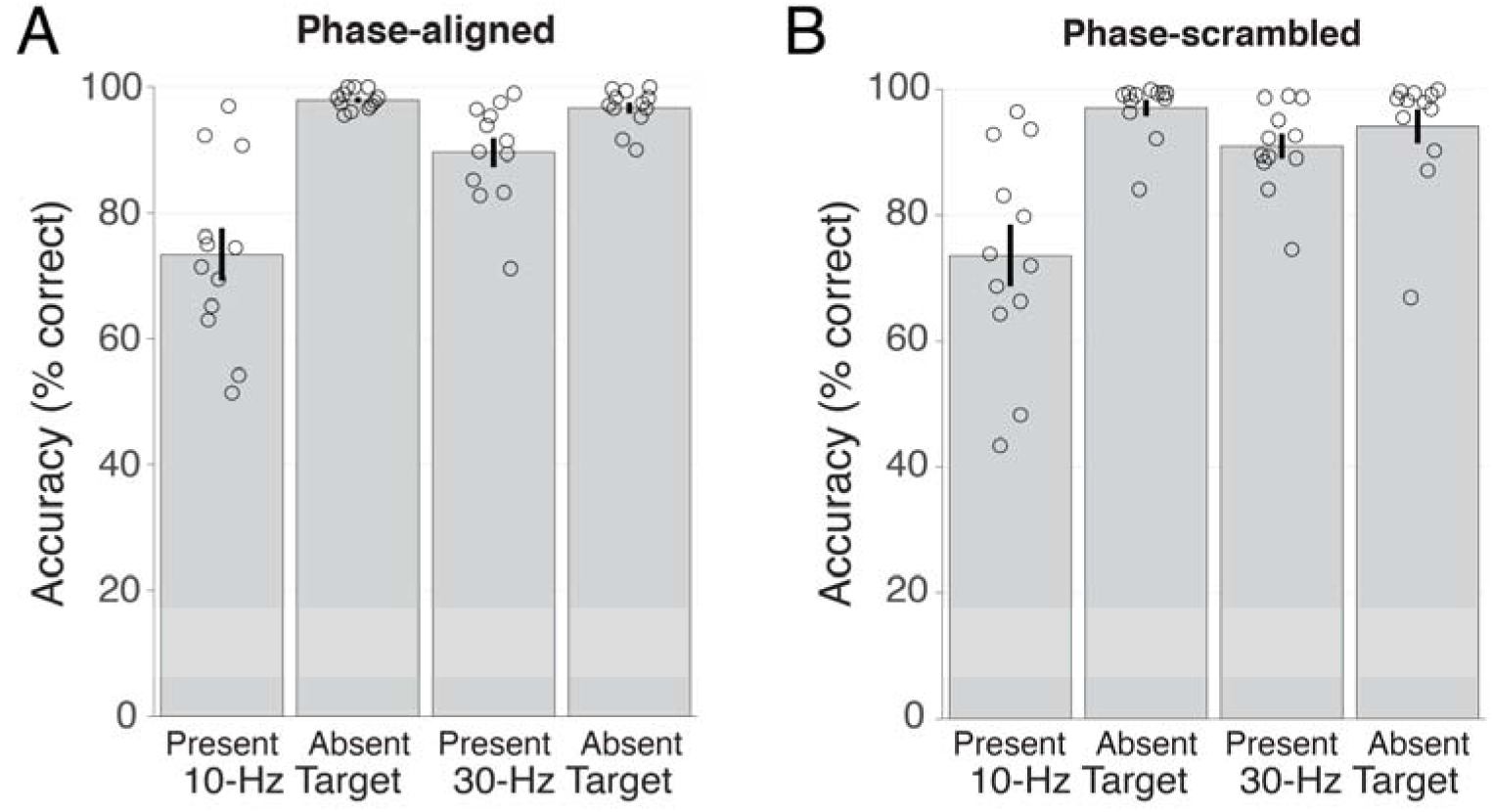
Performance across target frequency, target presence, and distractor phase conditions. (***A***) Mean accuracy under each condition with phase-aligned distractors. Open circles indicate individual subjects. Error bars represent s.e.m. (***B***) Mean accuracy under each condition with phase-scrambled distractors.

### Response times on tactile search trials

Figure 3A shows an example subject’s response times on correctly-performed search trials involving phase-aligned distractors. Four notable observations are clear from these data. First, response times were generally slower on trials when the target stimulus was absent compared to trials when the target stimulus was present. Second, response times tended to increase as a function of set size for both target-absent and target-present trials. Third, response times were slower when the subject searched for a 10-Hz target compared to searching for a 30-Hz target at the same set size. Finally, response time variations associated with set size differences were larger with the 10-Hz target trials compared to the 30-Hz target trials. These result patterns are also seen in the group-averaged data (Fig. 3B).

**Figure 3.**
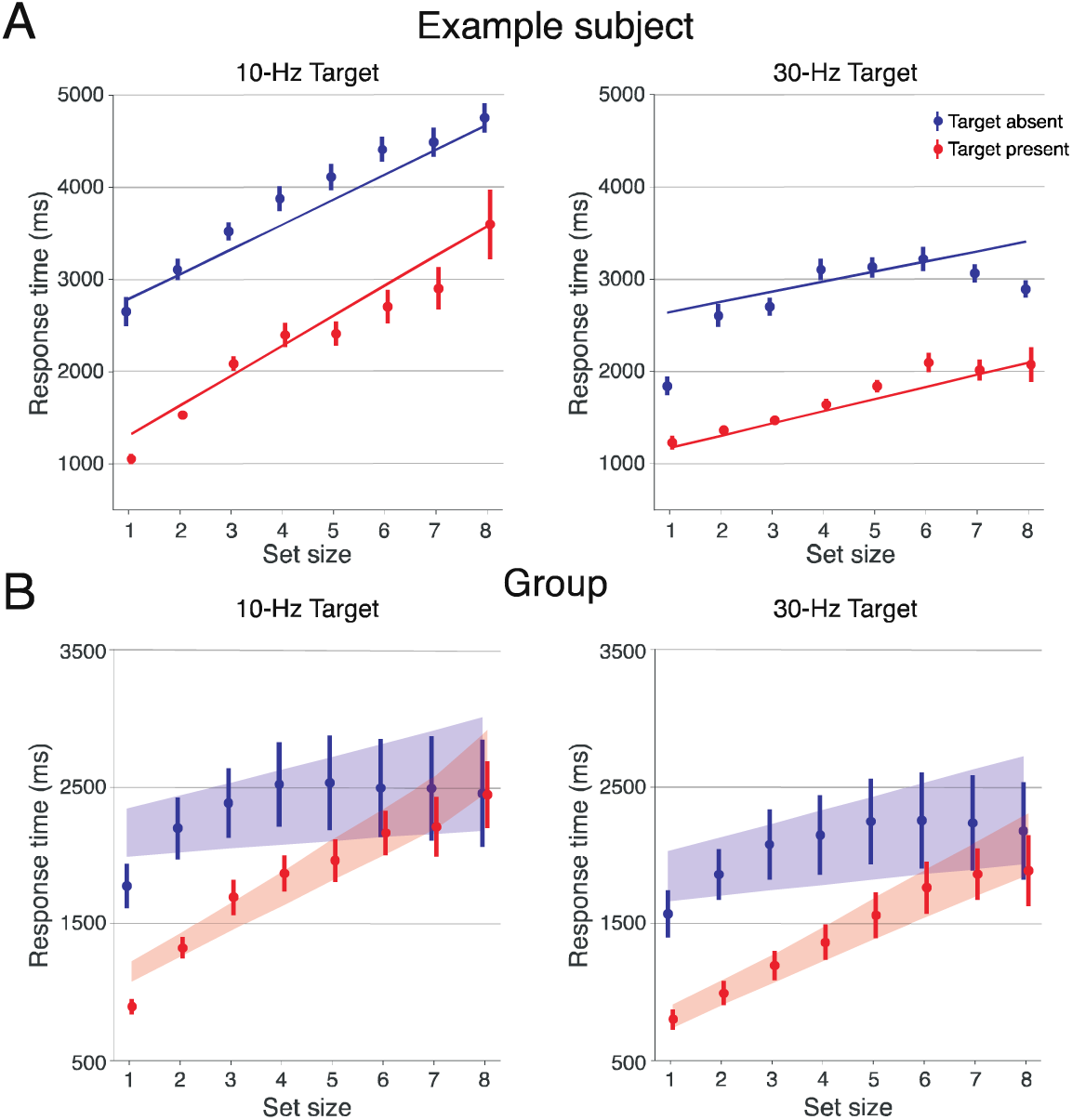
Tactile search times with phase-aligned distractors (N = 12). (***A***) Example subject results for correct trials only with 10-Hz (left) and 30-Hz targets (right). Mean response times for target-present (red) and target-absent (blue) trials as a function of set size. Error bars represent s.e.m. *(**B**)* Group results. Mean response times for target-present (red) and target-absent (blue) trials as a function of set size. Shaded regions indicate predictions from the linear mixed effects model (mean ± s.e.m.).

We statistically assessed the influence of target presence, target frequency, and set size using linear mixed effects modeling. Table 2 summarizes the model results for these experimental manipulations relative to performance achieved on target-present trials involving the 30-Hz target. Consistent with our qualitative assessments, these analyses revealed a significant effect of target presence (t_24207_ = 24.18, p = 1.31e-127) on overall response times, as the estimated intercept for the target-absent condition (1776 ms) was nearly 3-fold greater than the intercept for the target-present condition (653 ms). The observation that response times tended to be slower in the 10-Hz target condition compared to the 30-Hz target condition was also confirmed by a significant effect of target frequency on the intercepts (t_24207_ = 6.97, p = 3.31e-12). The effect of target presence did not depend meaningfully on target frequency as the target presence by target frequency interaction failed to achieve significance (p = 0.34). In addition to the effects on baseline response times (intercepts), the linear mixed effects model also provides slope estimates that captured the relationship between response time and set size. The slopes in all conditions were positive as response times increased with set size. In the visual search literature, slopes are used as a metric of search efficiency (Treisman and Gelade 1980) with lower slopes indicating more efficient search processes. With tactile search, slopes differed significantly according to target presence (t_24207_ = -10.51, p = 1.02e-25) as search appeared more efficient in the target-absent condition. Although the slope in the target-absent condition was 30-40% that of the target-present condition for both 10-Hz and 30-Hz targets, there were subtle slope differences that depended on target frequency, captured by a significant three-way interaction between set size, target absence, and target frequency (t_24207_= -2.79, p = 0.005) that was likely driven by the target-present condition where slopes were significantly larger for 10-Hz targets compared to 30-Hz targets (t_24207_= 3.43, p = 0.0006).

**Table 2.** Linear mixed effect model results for response times with phase-aligned distractors.

| **Fixed effects:** | **Estimate** | **S.E.M.** | **df** | **t value** | **p value** |
| --- | --- | --- | --- | --- | --- |
| *Baseline: 30-Hz target, target present* |  |  |  |  |  |
| Intercept | 652.39 | 143.21 | 13 | 4.56 | 0.0006 |
| Slope (set size) | 171.48 | 6.62 | 24207 | 25.91 | 5.31e-146 |
| *Intercept effects* |  |  |  |  |  |
| Target absent | 1123.42 | 46.47 | 24207 | 24.18 | 1.31e-127 |
| 10-Hz Target | 261.00 | 37.46 | 24207 | 6.97 | 3.31e-12 |
| 10-Hz Target:Target absent | 63.24 | 66.58 | 24207 | 0.95 | 0.34 |
| *Slope Effects* |  |  |  |  |  |
| Set size:Target absent | -104.78 | 9.98 | 24207 | -10.51 | 1.02e-25 |
| Set size:10-Hz Target | 35.03 | 10.22 | 24207 | 3.43 | 0.0006 |
| Set size:Target absent:10-Hz Target | -41.00 | 14.68 | 24207 | -2.79 | 0.005 |

Model fit to response times of correct trials only. Response times with the 30-Hz target, target-present condition are taken as the reference. Degrees of freedom (df), estimated using the Satterthwaite method, have been rounded to the nearest integer.

Our linear mixed model assumed a linear relationship between response times and set size, and this analysis approach generally captured the variance structure in our data; however, there were signatures of nonlinearity that were most obvious in the target-absent condition, particularly with 30-Hz targets. Indeed, in this condition, there was a clear inflection in the 30-Hz target response time profile of the example subject (Fig. 3A) with response times decreasing, rather than increasing, above set size 6. Notably, multiple subjects tested in this condition spontaneously reported experiencing a coherent percept of the 10-Hz distractors over the body at larger set sizes. We hypothesized that this subjective experience, and the facilitated performance at large set sizes with the 30-Hz target in the target-absent condition, may have reflected perceptual grouping of the phase-aligned 10-Hz distractors. We reasoned that scrambling the phase of the distractor stimulus trains (Fig. 1B) would disrupt their coherence and prevent grouping. Performance patterns with phase-scrambled distractors (Fig. 4A) were mostly consistent with the performance observed with phase-aligned distractors. The linear mixed model results for phase-scrambled distractors are summarized in Table 3. Again, we found that response times were significantly faster on target-present trials compared to target-absent trials (i.e., target presence effect on intercepts; t_24214_ = 18.42, p = 2.79e-75). Response times were again significantly faster with the 30-Hz targets compared to 10-Hz targets (target frequency effect on intercepts: t_24214_ = 10.71, p = 1.04e-26). Notably, because the difference in intercepts between the target-present and target-absent conditions were substantially larger with 10-Hz targets compared to 30-Hz targets, we also observed a significant target presence X target frequency interaction (t_24214_ = 4.78, p = 1.81e-06). While this interaction effect, which was not observed with the phase-aligned distractors, could simply reflect a net response time reduction with 30-Hz targets in the phase-scrambled conditions, it is also consistent with changes in the linearity of the response time profiles and a better correspondence between the predictions of the linear model at the small set sizes.

**Figure 4.**
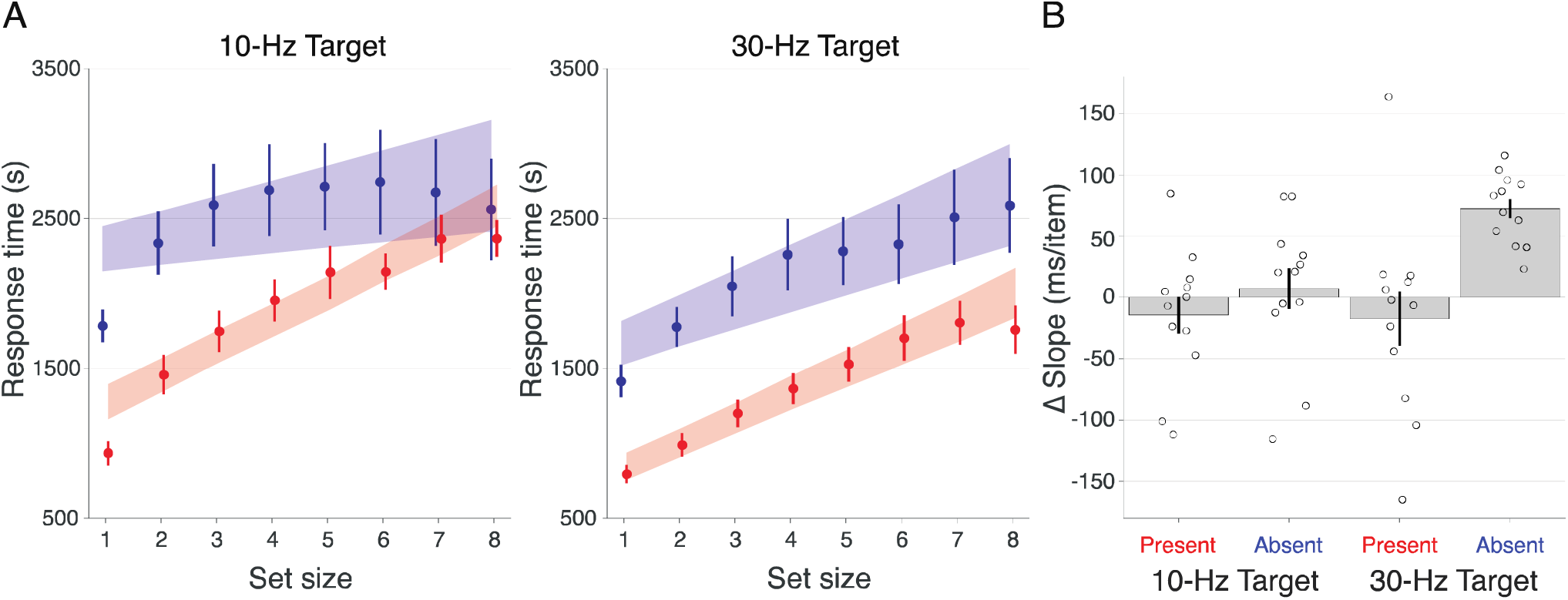
Tactile search times with phase-scrambled distractors (N = 12). (***A***) Group results showing mean response times for target-present (red) and target-absent (blue) trials as a function of set size. Conventions as in Fig. 3B. (***B***) Bar plot depicts the slope differences between the phase-aligned and phase-scrambled conditions (scrambled – aligned). Open circles indicate individual subjects. Error bars represent s.e.m. Consistent and significant (*p* < 0.05) slope changes are only observed with the 30-Hz target, target-absent condition.

**Table 3.** Linear mixed effect model results for response times with phase-scrambled distractors.

| **Fixed effects:** | **Estimate** | **S.E.M.** | **df** | **t value** | **p value** |
| --- | --- | --- | --- | --- | --- |
| *Baseline: 30-Hz target, target present* |  |  |  |  |  |
| Intercept | 693.00 | 105.25 | 13 | 6.58 | 1.60e-05 |
| Slope (set size) | 157.55 | 6.16 | 24214 | 25.57 | 2.84e-142 |
| *Intercept effects* |  |  |  |  |  |
| Target absent | 830.58 | 45.08 | 24214 | 18.42 | 2.79e-75 |
| 10-Hz Target | 383.88 | 35.84 | 24214 | 10.71 | 1.04e-26 |
| 10-Hz Target:Target absent | 308.63 | 64.64 | 24214 | 4.78 | 1.81e-06 |
| *Slope Effects* |  |  |  |  |  |
| Set size:Target absent | -15.96 | 9.58 | 24214 | -1.67 | 0.10 |
| Set size:10-Hz Target | 17.79 | 9.59 | 24214 | 1.86 | 0.06 |
| Set size:Target absent:10-Hz Target | -89.12 | 14.08 | 24214 | -6.33 | 2.46e-10 |

Model fit to response times of correct trials only. Response times with the 30-Hz target, target-present condition are taken as the reference. Degrees of freedom (df), estimated using the Satterthwaite method, have been rounded to the nearest integer.

Analysis of slope estimates also revealed notable differences between performance achieved with phase-scrambled distractors compared to phase-aligned distractors. Although we again observed a significant three-way interaction between set size, target presence, and target frequency (t_24214_ = - 6.33, p = 2.46e-10), the robust main effect of target presence observed with phase-aligned distractors was no longer significant. This is likely due to the fact that, with phase scrambling, slope differences between target-absent and target-present conditions became reduced with 30-Hz targets (i.e., the slope in the target-absent conditions was ∼90% that of the target-present conditions) while slope differences remained large with 10-Hz targets (i.e., the slope in the target-absent conditions is ∼40% that of the target-present conditions). Importantly, this change in the slope differences appeared to be attributable to slope changes in the target-absent condition with the 10-Hz distractors rather than the target-present condition. Moreover, the increased slopes with phase scrambling in the target-absent condition is consistent with longer response times at the large set sizes with the 10-Hz distractors, the trial types that appeared most vulnerable to potential distractor grouping effects.

To compare slopes achieved under the phase-aligned and phase-scrambled distractor conditions more explicitly, we fit linear functions to the averaged response times sorted by set size at a single-subject level separately for the target presence X target frequency conditions (i.e., for each subject, we estimated 8 slopes; 2 target presence conditions X 2 target frequency conditions X 2 distractor phase conditions). For each of the 4 target presence X target frequency conditions, we then determined how slopes differed between the phase-scrambled and phase-aligned conditions (Fig. 4B). The only condition where we saw a consistent change in slope related to phase scrambling was the target-absent condition with 30-Hz targets (involving the 10-Hz distractors), where larger slopes were observed in the phase-scrambled condition compared to the phase-aligned condition in all 12 subjects. The average slope in this condition was significant according to a one-sample t-test against the null hypothesis that slopes were unchanged between phase conditions (t_11_ = 8.812, p = 2.58e-06).

### Response time contributions of individual body-sites

To characterize how tactile stimulation of a single body-site affected search times, we fit a linear model describing each location’s response time contributions that allowed for distinct target and distractor effects. These simple linear models generally accounted for response time variations across the target frequency and distractor phase conditions (root mean square error (RMSE) = 0.81 ± 0.05 ms). Because RMSE did not differ significantly according to condition (F(3,44) = 0.55, p = 0.65), we averaged the target and distractor weights across conditions (Fig. 5). Average weights for the target terms tended to be negative, indicating that the presence of targets was generally associated with faster response times relative to baseline (Fig. 5A). While average target weights did not differ significantly according to body-site (F(7,88) = 1.53, p = 0.17), the target weights associated with the index fingers generally were of the greatest absolute magnitude. This pattern may account for the finding that target response times differ significantly according to body-site (F(7,88) = 5.7, p = 1.87e-05) with the fastest responses occurring with index finger targets. The uniqueness of the index fingers was also revealed by the average distractor weights (Fig. 5B). These weights tended to be positive indicating that distractors on a single-site generally increased response times relative to baseline. As with target weights, the distractor weights with the greatest magnitude were associated with the index fingers. With distractor weights, significant differences related to body-site were observed (F(7,88) = 2.47, p = 0.02), and these site-dependent differences failed to achieve significance when index finger weights were omitted (F(5,66) = 0.6, p = 0.69). Site-dependent distractor weights can explain the site-dependent response times for single-site distractor trials (F(7,88) = 2.65, p = 0.016).

**Figure 5.**
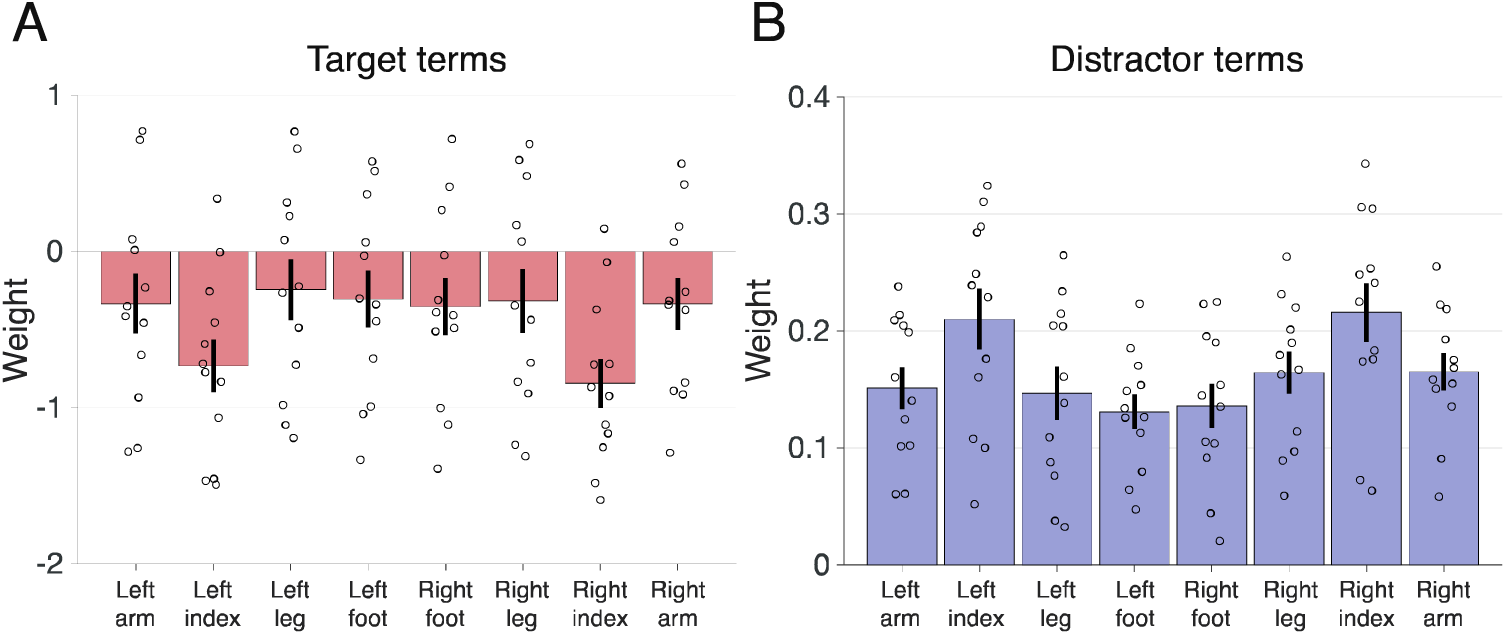
Single-site model weights. (***A***) Mean weights for each target site averaged over conditions and subjects. (***B***) Mean weights for each distractor site averaged over conditions and subjects. Open circles indicate individual subjects. Error bars represent s.e.m.

### Response time contributions of 2-site combinations

Although the single-site models explained a substantial amount of response variance, we reasoned that performance on trials involving multiple co-stimulated sites may have reflected interactions between simultaneously stimulated body locations rather than merely the independent contributions of isolated sites. To test this possibility, we used randomized matching pursuit (Materials and Methods) to identify pairs of stimulated sites that, in conjunction, may have differentially influenced response times. In these models, we assumed that two-site terms contributed to responses times in addition to the single-site target and distractor terms. These models, like the single-site models, ignore explicit effects of size set, which can be independent of the spatial location of the tactile stimuli. For each condition in each subject separately, we used a rigorous model selection procedure to determine the number of two-site terms to include in the model, resulting in 48 fitted models (Fig. 6). This procedure, based on BIC, provided an objective way to balance model complexity with improvements in model performance as indexed by reductions in prediction error (Fig. 6A). Nearly all models (47/48) included at least one two-site term with an average of 36.42 ± 2.5 two-site terms included in the BIC-preferred models. Although model performance (RMSE) did not differ significantly according to target frequency or distractor phase (main and interaction effects, all ps > 0.29), model complexity differences were observed (Fig. 6B). The number of two-site terms differed significantly according to distractor phase (main effect: F(1,44) = 4.2, p = 0.046) with more terms included in models of the phase-aligned data. The number of two-site terms did not depend on target frequency (main effect: F(1,44) = 0.005, p = 0.95; interaction effect: F(1,44) = 0.19, p = 0.67).

**Figure 6.**
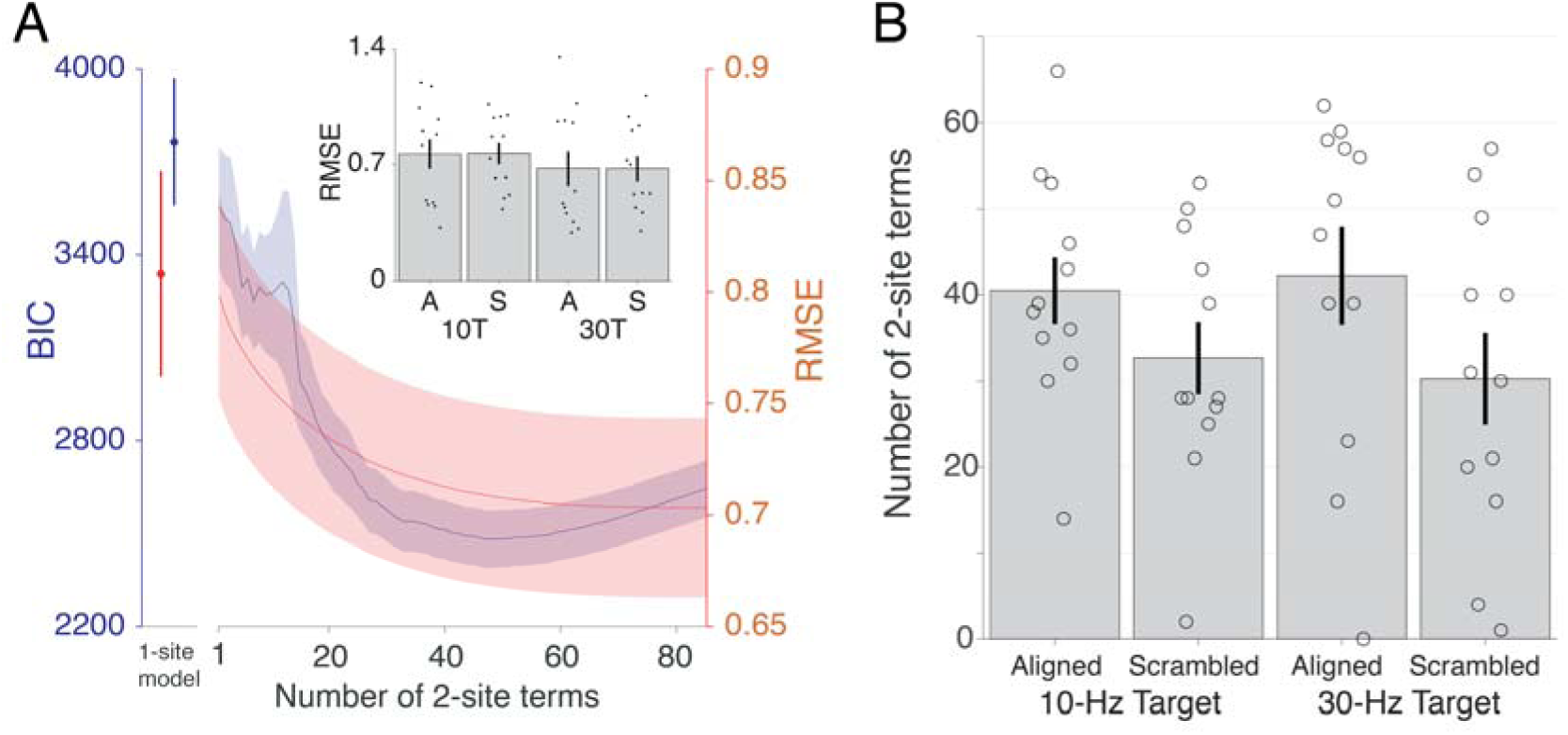
Two-site model complexity and performance. (***A***) Blue trace depicts that group-averaged Bayesian information criterion (BIC) for 2-site models as a function of included number of 2-site terms. Shaded region indicates s.e.m. Blue marker indicates group-averaged BIC for 1-site models. Error bar represents s.e.m. Red trace depicts that group-averaged model performance, indexed by root mean square error (RMSE) for 2-site models as a function of included number of 2-site terms. Shaded region indicates s.e.m. Red marker indicates group-averaged RMSE for 1-site models. Error bar represents s.e.m. Inset bar plot depicts condition-specific average RMSE for BIC-preferred models for 10-Hz and 30-Hz targets with phase-aligned distractors (A) and phase-scrambled distractors (S). Dots indicate individual subjects. (***B***) Bar plot depicts the average number of two-site terms included in the BIC-preferred model for each condition. Open circles represent individual subjects. Error bars indicated s.e.m.

Despite the variable number of two-site terms across subjects and conditions (Fig. 6B), a substantial number of two-site terms – both target-distractor and distractor-distractor pairs – were consistently included in the BIC-preferred models (Fig. 7). In fact, a given two-site term was included in 24.77 ± 0.69% of the models on average and 22 terms appeared at least once in a model in all 12 subjects. We used Hamming distance to quantify the similarity of BIC-preferred models (Materials and Methods). Within subjects, we compared model similarity between the 10-Hz and 30-Hz conditions, with terms concatenated over phase conditions, and found that the average Hamming distance (0.478 ± 0.26) was significantly lower (i.e., more similar) than expected by chance (randomization test, p < 0.005). We also compared model similarity between pairs of subjects, concatenating over target frequencies and phase conditions: The average between-subject Hamming distances was 0.476 ± 0.01, which was significantly lower than expected by chance (randomization test, p < 0.005). These results indicate that randomized matching pursuit and model selection identified consistent two-site terms both within and between subjects.

**Figure 7.**
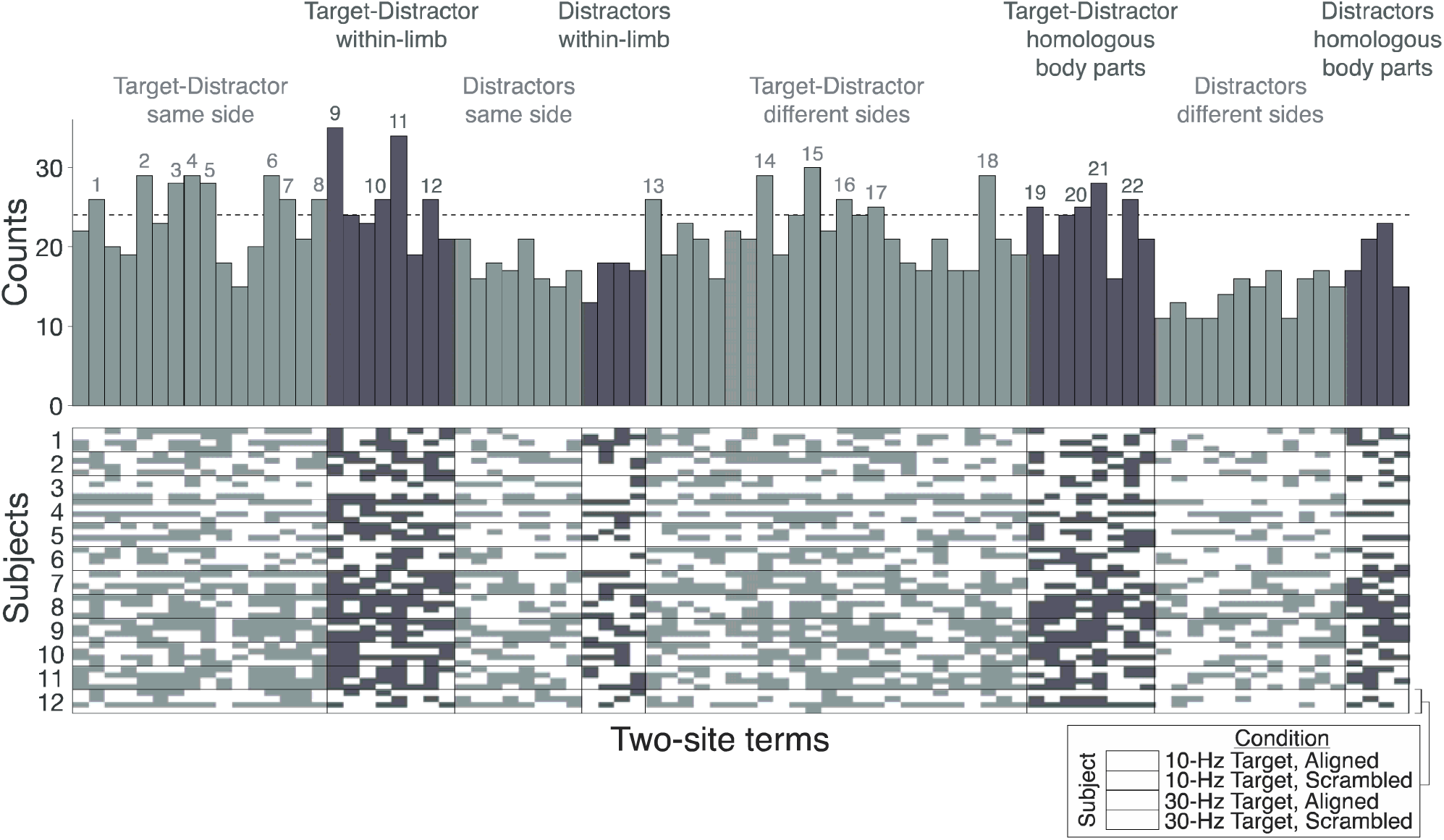
Distribution of two-site terms observed in BIC-preferred models. Histogram (top panel) indicates frequency of individual two-site terms over all fitted models. Terms are grouped by category. Numbered terms are those which appeared in more than 50% of models (dashed line indicates threshold; see Fig. 8). The presence of each two-site term in each subject’s 4 models are depicted in the binarized vector plots (bottom panel).

To establish whether interactions between particular pairs of body-sites exerted greater influences on tactile search times, we focused on the two-site terms that appeared most frequently in preferred models. Twenty-two terms occurred in greater than 50% of the models (Fig. 8). These only comprised target-distractor combinations and nearly all (21/22) resulted in response time increases relative to baseline. The target-distractor terms could be grouped into 4 categories: within-limb, ipsilateral (different limbs, same side), homologous (same site, different sides), or contralateral (non-homologous). These patterns reveal a modest structure in two-site interactions that are defined according to both side and body part. Notably, the average weight of the terms involving ipsilateral stimulation sites (0.36 ± 0.05) were significantly higher than that of the terms involving contralateral stimulation sites (0.15 ± 0.04) (t_20_ = 3.25, p = 0.004), implying that stimulation of two sites on the same side of the body, particularly within the same limb, preferentially modulated tactile search times.

**Figure 8.**
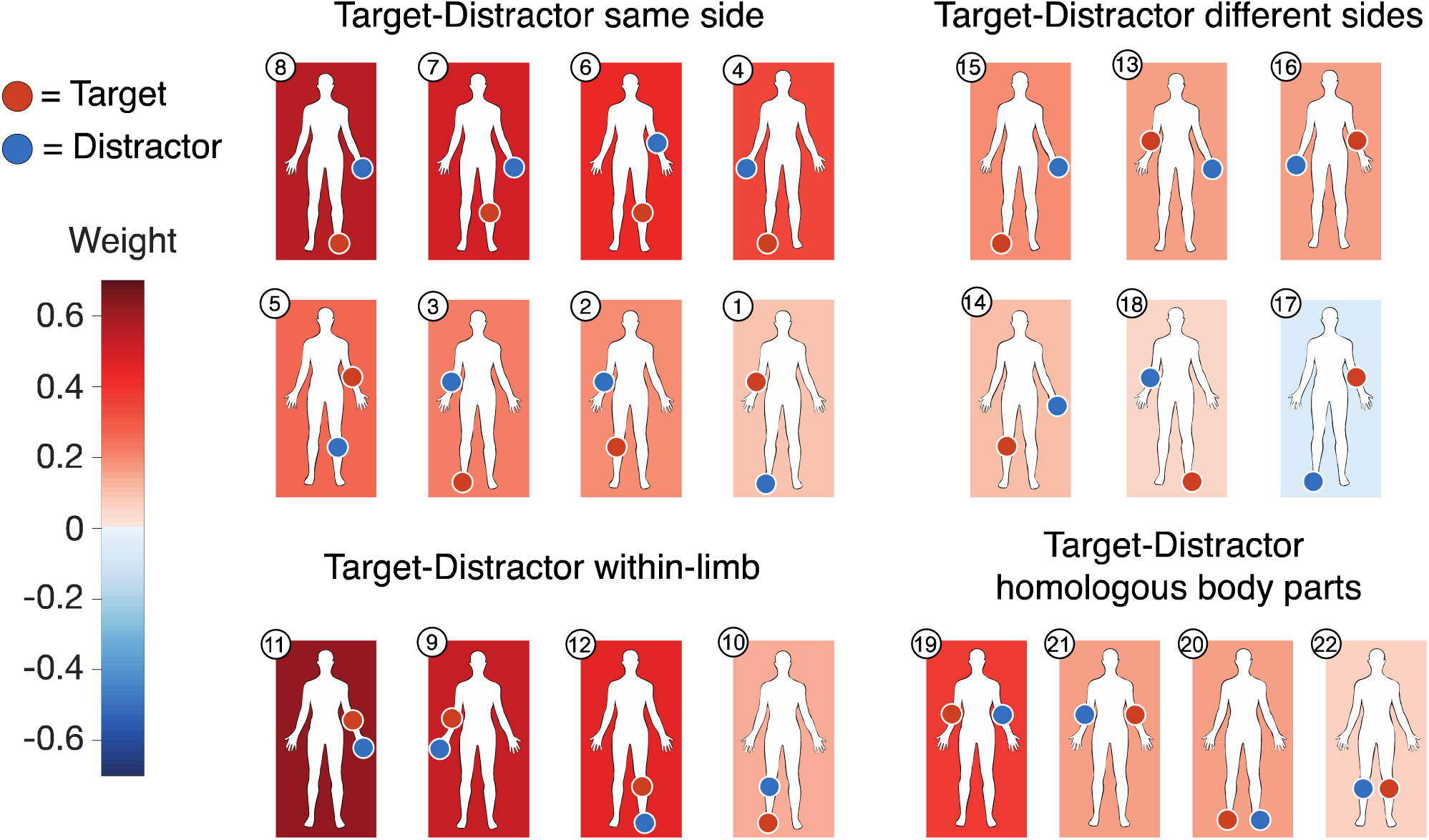
Two-site terms that occurred most frequently across sessions. Twenty-two terms identified as occurring in more than 50% of BIC-preferred models (Fig. 7). Each panel depicts location of target (red circle) or distractor (blue circle). Background color indicates the term’s mean weight across sessions. Terms are grouped by category.

## Discussion

We asked participants to report the detection of target tactile stimulation over their body in the absence of distractor cues and in the presence of a variable number of distractors. Participants generally performed the task accurately and we characterized response time variations to infer how tactile search performance related to target presence, set size, target frequency, and stimulation sites. We observed longer response times on distractor-only trials compared to trials comprising a target. With target-present and target-absent trials, response times typically increased with the total number of stimulated sites. Response times and search slopes differed depending on the target frequency (10Hz vs 30Hz) so tactile search of flutter trains can be asymmetric. Some of this asymmetry may have been due to perceptual grouping of distractors, which was more pronounced with 10-Hz distractor trains. We modeled the response time contributions of individual body-sites and found that search behavior was more influenced by stimulation on the index fingers compared to other sites. We also modeled the modulatory effects of body-site pairs and found that co-stimulation of ipsilateral sites, particularly those within a limb, most consistently influenced tactile search times.

With visual search, positive slopes in the linear functions relating search time with set size are taken as evidence for inefficient search and a need to serially process items in the display (Treisman and Gelade 1980). Because we found that response times nearly always increased with set size, we infer that full-body search for tactile cues in the flutter range is largely a serial process. Indeed, we observed response time increases of ∼60-70 ms/item on target-absent trials and ∼180-210 ms/item on target-present trials, with subtle differences depending on the target frequency (see below). These slopes are far larger than those typically encountered in visual search, where slopes of 10-30 ms/item reflect inefficient search (Duncan and Humphreys 1989; Eckstein 2011). Importantly, tactile search slopes can be in the lower ranges occupied by visual slopes (Lederman and Klatzky 1997; Overvliet et al. 2007; Toet et al. 2008) and larger than the values we observed (Lederman and Klatzky 1997), depending on the target features and stimulated locations, so there may not be simple modality-dependent slope differences.

We found that search times tended to be longer on trials when no target was presented, consistent with other tactile search data (Lederman and Klatzky 1997; Overvliet et al. 2007; Toet et al. 2008) and visual search behavior (Duncan and Humphreys 1989; Treisman and Gelade 1980). This pattern makes sense assuming serial search because, for a given set size, participants would need to evaluate more sites on target-absent trials compared to target-present trials before determining their response thus requiring more time on average. However, this logic would also predict that search slopes would be steeper in the target-absent condition compared to the target-present condition, as often seen in visual search tasks (Eckstein 2011; Wolfe and Horowitz 2017). We observed the opposite pattern: Slopes for the target-absent condition were consistently lower than slopes for the target-present conditions (Figs. 3B and 4A). The lower slopes in the target-absent conditions, which have also been reported in other tactile search paradigms (Lederman and Klatzky 1997; Overvliet et al. 2007; Toet et al. 2008), may be related to perceptual grouping effects that appear to be more pronounced in the absence of target cues. Indeed, using phase-aligned distractors, response time profiles deviated from linear model predictions, particularly at larger set sizes with the 10-Hz distractors. Notably, in these conditions, participants subjectively reported that they experienced the 10-Hz trains as coherent sensations over their body. Conceivably the grouping of multiple 10-Hz distractors based on their temporal coherence facilitated search: Participants could have decided on the presence or absence of the target cue simply by judging whether they experienced a signal that “popped out” from the coherent distractor pattern. This possibility is supported by the observation that disrupting the phase relationship between distractors systematically increased search slopes with the 10-Hz distractors (Fig. 4B). It is important to consider why these effects depended on target (or distractor) frequency. These frequency-dependent search asymmetries – differences when the target and distractor assignments of the 10- and 30-Hz stimulation is swapped – are likely attributable to tactile temporal binding windows and thresholds for timing differences between multiple tactile cues. In fact, tactile simultaneity judgment (TSJ) thresholds are ∼60-70 ms (Heed and Azañón 2014; Shore et al. 2005; Yamamoto and Kitazawa 2001). With phase-aligned 10-Hz distractors, inter-stimulus intervals experienced over the body were either 0 or 100ms, which likely facilitated the perception of simultaneously co-occurring events (over sites) and discrete events (between cycles) thereby promoting grouping. With phase-scrambled 10-Hz distractors, as with 30-Hz distractors in both phase conditions, inter-stimulus intervals fell under the TSJ threshold thereby obscuring the relationships between the distractor trains and weakening grouping effects. Our results thus imply that timing information can serve to bind tactile cues, similar to reported effects in vision (Cass et al. 2011) and audition (Bregman 2009). Assuming that timing cues are solely responsible for the perceptual grouping of vibrations (which is almost certainly not the case), we would predict less asymmetry if 50-Hz stimulation was tested with 30-Hz rather than 10-Hz stimulation. Additional experiments are required to better understand frequency-dependent effects in tactile search over the body.

We explored the contributions of individual body sites and body-site pairs on tactile search times. When stimulation sites were assumed only to have independent effects, with distinct target and distractor influences, we found that stimulation on the index fingers exerted greater modulatory effects on response times. This result is notable because we matched the stimulation amplitudes across all sites and target frequencies for perceived intensity. Accordingly, the results from the independent-site models cannot be trivially explained by differences in the perceived intensities of the stimuli, as least as measured when the cues were delivered in isolation (Materials and Methods). Instead, the relatively large weights associated with stimulation of the index fingers likely reflect the dominance of the index fingers in multi-site interactions or masking effects (Gilson 1969). This point is also supported by the frequent involvement of the index fingers in the two-site model terms identified by randomized matching pursuit and model comparison. These models, which capture pairwise interactions between body-sites, also show that co-stimulation of sites within the same limb or on the same side of the body tend to have greater influences on search times. This result pattern may also be related to principles underlying masking in the somatosensory system, as earlier studies have reported stronger masking between sites that are closer in somatotopic space (Gilson 1969) and stronger masking with ipsilateral compared to contralateral stimulus patterns (Levin and Benton 1973; Sherrick 1964). Interestingly, masking effects between contralateral cues tend to be stronger between homologous body parts (D’Amour and Harris 2014b) and our models also revealed the consistent contribution of these two-site terms. The spatial principles underlying our data, irrespective of how they relate to masking, appear to reflect the lateralized representations of the body that exist in multiple levels of the somatosensory neuraxis and perhaps indicate how attention signals operate on somatotopic maps. Crucially, our results are also consistent with spatial processing that operates in external space rather than a body-based coordinate system (Heed and Azañón 2014; Medina and Coslett 2010; Röder et al. 2002). Indeed, multi-site tactile interactions can be modulated by the locations of the stimulated sites in space (Farnè et al. 2008; Rahman and Yau 2019) and cued attentional deployment over the body also appears to operate in external space rather than a simple body-based reference frame (Lakatos and Shepard 1997). Thus, although we found consistent spatial patterns, additional experiments that manipulate posture are required to establish a clearer understanding of the spatial principles underlying tactile search behavior.

There are a number of study limitations to be noted. First, because we tested each subject so extensively, this necessarily required many test sessions. By carefully randomizing and distributing the conditions within each session and counter-balancing session orders and conditions across subjects, we ensured that our specific result patterns could not be simply attributable to general learning effects that may have been introduced with repeated testing. Although there was a trend for response times to reduce over the 8 test sessions, this trend was not significant (F(7,88) = 0.5, p = 0.8307). Related to this, although we carefully established each participant’s single-site stimulus amplitudes, equating all by perceived intensity, at the outset of the experiments, we did not retest these amplitudes to confirm consistency over time. Second, we exerted minimal control on where or how participants deployed their spatial attention to their body, although we required most participants to maintain central visual fixation during testing (Materials and Methods). This is an important point because search times, especially in serial processes, likely depend on the starting locus of attention and there is evidence that the perception of complex tactile patterns over the body depends on how attention is spatially cued (Lakatos and Shepard 1997). We predict that the variance of measured response times would be reduced if attention is more systematically manipulated, but this needs to be tested explicitly. Finally, a critical aspect of our experimental design worth consideration is the fact that subjects quickly learned that there was a finite number of locations on which tactile stimulation could be experienced (i.e., the locations where tactors were placed). This differs from the conditions of visual search paradigms, where target and distractor stimuli can appear essentially in a far larger number of configurations and positions that more continuously span the visual space (i.e., that of a monitor). Given subjects’ knowledge of the restricted but fixed stimulation sites, they conceivably could adopt particular strategy for monitoring and searching based on those sites and could change if tested in another context or if they lack this knowledge. Ongoing studies in the lab are focused on this important consideration.

We investigated tactile search over the body, focusing on the different limbs rather than on a single limb or body “part” (De Vignemont et al. 2009), in an effort to avoid potential confounds related to the physical interactions between stimuli or more pronounced masking effects. There have been extensive efforts to characterize tactile search behavior using analogs of the visual search task, but these have nearly always focused on the processing of tactile stimulation on the hands (Lederman et al. 1988; Lederman and Klatzky 1997; Overvliet et al. 2007) or a single body part like the torso (Toet et al. 2008). These studies also characterized search behavior using a multitude of stimulus features covering spatial domains (e.g., oriented bars, edges, shapes), temporal domains (e.g., duration), and surface properties (e.g., roughness, compliance, temperature). To our knowledge, few studies have quantified tactile search behavior using stimuli similar to ours (although see Forster et al. 2016). Our results are consistent with some earlier findings: Response times tend to be longer on target-absent trials compared to target-present trials and response times, in many cases, increase with set size indicating serial processing. However, our results also differ from earlier reports: The relationship between search times and set size can vary dramatically, parallel processing appears to be possible in some cases, and search need not be asymmetric. These myriad results imply that there are many differences in search behavior that depend on the specific stimulus features. That said, more comprehensive testing and analyses that systematically compare and contrast search behavior achieved with different features may still reveal general principles for tactile search like the guiding features that explain visual search (Wolfe and Horowitz 2017). Establishing such principles may be important for the study and treatment of body schema disorders, such as phantom limb and body dysmorphia (Haggard and Wolpert 2004), and in commercial and industrial applications that use simultaneous touch for haptics-based communication (van Erp et al. 2005; Furlanetto et al. 2014; Gilson et al. 2007; Hagelsteen et al. 2019).

## Acknowledgements

This work was performed in the Neuromodulation and Behavioral Testing Facilities of BCM’s Core for Advanced MRI (CAMRI). We thank Yau Lab members for helpful discussions. This work was supported by BCM seed funds and the Alfred P. Sloan Research Fellowship.

